# Peptide allosteric inhibitor of TNFR1 signaling attenuates inflammation and rheumatoid arthritis pathology in human TNF transgenic mice

**DOI:** 10.64898/2026.06.30.735704

**Authors:** Tanver Hasan Riyed, Kriti Kalary, Jialiu Zeng, Chih Hung Lo

**Affiliations:** Department of Biomedical and Chemical Engineering, Syracuse University, Syracuse, NY 13244; Department of Biology, Syracuse University, Syracuse, NY 13244

**Author notes:** Correspondence: Chih Hung Lo, PhD.

**Keywords:** Therapeutic peptide, allosteric inhibitor, TNFR1 signaling, chronic inflammation, rheumatoid arthritis, transgenic mice

## Abstract

Inhibition of tumor necrosis factor receptor 1 (TNFR1) represents a major therapeutic strategy for chronic autoimmune and inflammatory diseases such as rheumatoid arthritis (RA). As current anti-TNF therapies can cause adverse side effects due to global blockade of the ligand, receptor-specific inhibition of TNFR1 signaling has emerged as a highly sought-after strategy. We have recently identified a novel peptide-based allosteric inhibitor, FKC (FKCRRWQWRMKK), that targets TNFR1 conformationally active region to alter receptor conformational states and disable receptor-ligand signaling complex. Here, we evaluated the therapeutic efficacy of FKC in a human TNF (hTNF) transgenic mouse model of RA. FKC treatment improves clinical RA scores in hTNF mice, accompanied by enhanced grip strength and increased walking distance. Importantly, FKC treatment inhibits TNF/TNFR1-mediated inflammation and attenuates RA pathology in hTNF mice. Together, our findings establish FKC as a promising new class of peptide-based therapeutics for chronic inflammatory diseases through selective inhibition of TNFR1 signaling.

## Introduction

Rheumatoid arthritis (RA) is a chronic autoimmune and inflammatory disease afflicting more than 18 million people worldwide [1]. It is characterized by progressive synovial inflammation, cartilage degradation, and periarticular bone erosion that ultimately culminates in irreversible joint destruction and functional disability [2]. The pathological hallmarks of RA joints include synovial hyperplasia with formation of an invasive pannus, infiltration of innate and adaptive immune cells including macrophages and lymphocytes, proteoglycan loss from articular cartilage, and osteoclast-mediated subchondral bone damage [3]. Collectively, these processes are orchestrated by a dysregulated cytokine network in which tumor necrosis factor (TNF) plays a central pathogenic role in mediating and propagating inflammation in RA [4].

TNF drives RA pathology primarily through TNF receptor 1 (TNFR1), which leads to IĸBα degradation and activates canonical NF-κB signaling to induce pro-inflammatory cytokines, chemokines, and matrix metalloproteinases that sustain synovial inflammation and joint destruction [5]. TNF is abundantly produced by activated macrophages and infiltrating immune cells in the arthritic synovium, with elevated levels correlating with disease severity and progression [5]. Although anti-TNF biologics have revolutionized RA treatment, global TNF blockade suppresses both TNFR1-driven inflammatory pathway and TNFR2-mediated immune regulation, leading to adverse side effects including increased infection risk and tuberculosis reactivation [6]. In addition, up to 40% of patients exhibit inadequate responses to initial anti-TNF therapy [7]. These challenges have positioned selective inhibition of TNFR1 signaling as an emerging next-generation therapeutic strategy for RA treatment [8].

Among TNFR1 receptor-specific targeting strategies, it has been demonstrated that disruption of receptor-ligand [9,10] and receptor-receptor interactions via competitive inhibition [11–13] as well as perturbation of receptor conformational dynamics via non-competitive inhibition or allosteric modulation [14–17] by small molecules and peptides are both feasible approaches to selectively modulate TNFR1 signaling [18]. Specifically, allosteric inhibitors that neither interfere with ligand binding nor compete with receptor-receptor interactions have been proposed to be more efficient because they are unencumbered by competition [14,19]. Importantly, a recent discovery of the inter-monomeric space between TNFR1 dimers as a key binding pocket for allosteric modulators [16] has led to the development of FKC (FKCRRWQWRMKK) peptide as a novel allosteric inhibitor of TNFR1 signaling [17]. FKC inhibits TNFR1 signaling by targeting the receptor conformationally active region, altering receptor conformational states, and disabling receptor-ligand signaling complex, without ablating receptor-ligand or receptor-receptor interactions (**Fig. 1A**) [17]. These findings support the emerging concept that receptor conformational state functions as a critical molecular switch in controlling receptor signaling [20,21].

**Figure 1.**
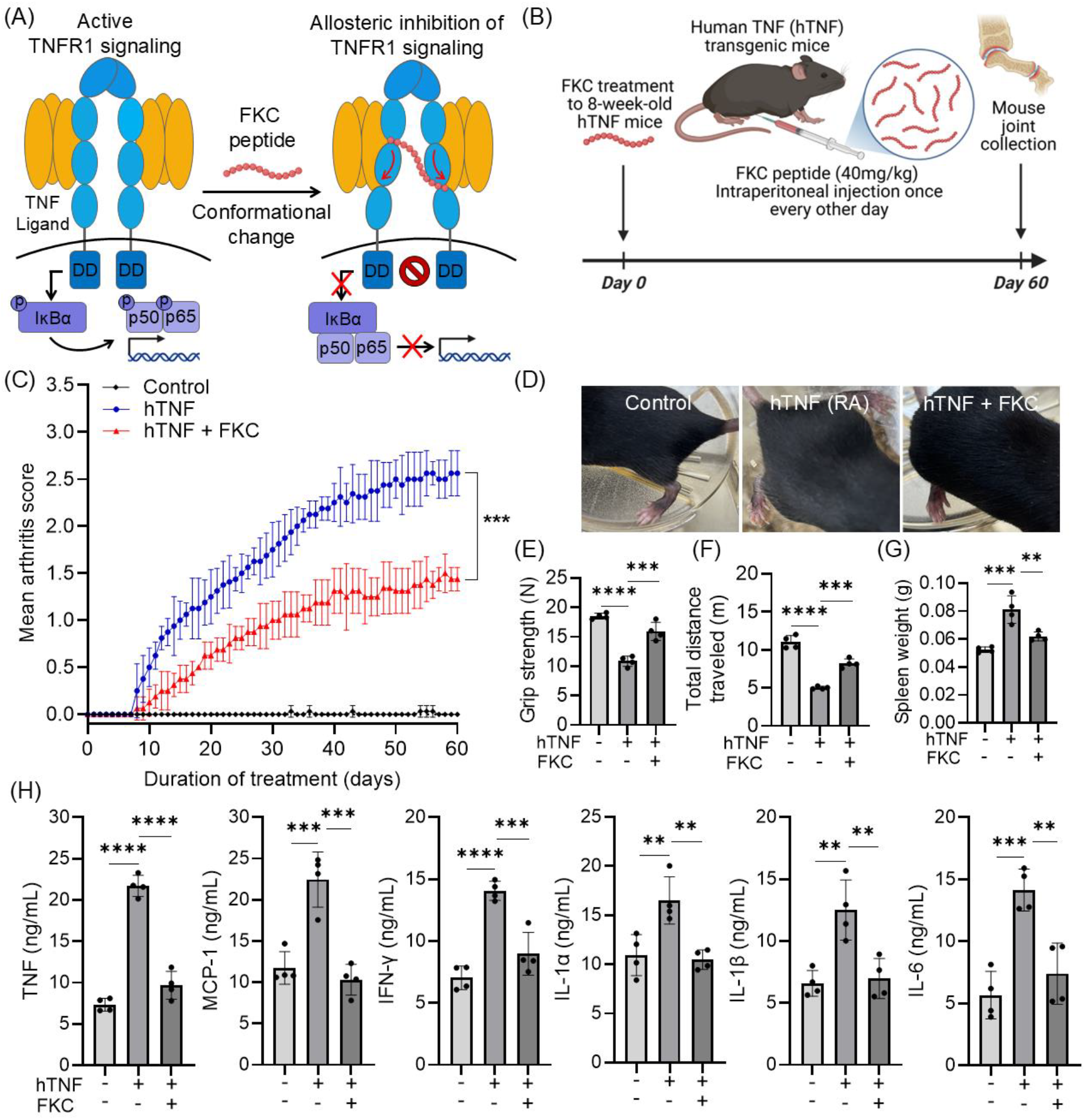
(A) Schematic illustrating the allosteric inhibition mechanism of FKC. FKC binds to TNFR1 and perturbs its conformational dynamics without disrupting receptor-ligand or receptor-receptor interactions. Upon FKC binding, the receptor dimer adopts an open, inactive conformation that is unable to recruit downstream signaling molecules, resulting in nonfunctional receptor-ligand complexes that cannot propagate signaling. (B) Schematic of the treatment regimen in hTNF mice. Eight-week-old hTNF mice are treated with FKC (40 mg/kg) via intraperitoneal injection every other day for 60 days. (C) Clinical scores monitored throughout the treatment period show that FKC reduces disease severity in hTNF mice. (D) Representative images of joint swelling in hTNF mice, demonstrating attenuation following FKC treatment. (E) Hindlimb grip strength measurements, indicating improved muscle function in FKC-treated mice. (F) Total distance traveled in the open field maze (OFM) across different groups, showing enhanced mobility following FKC administration. (G) Spleen weight measurements across treatment groups, with FKC-treated mice exhibiting reduced spleen size, consistent with decreased systemic inflammation. (H) In hTNF mice, plasma levels of pro-inflammatory cytokines (TNF, MCP-1, IFN-γ, IL-1α, IL-1β, and IL-6) are elevated, and FKC treatment reduces all measured cytokines. Data are presented as mean ± SD (N = 4 mice). **P < 0.01, ***P < 0.001, ****P < 0.0001 by one-way ANOVA with post hoc Tukey’s test for multiple comparisons.

While FKC has been shown to attenuate acute systemic inflammation in mice [17], its functional efficacy has not yet been examined in the context of chronic inflammation such as in RA, which differs substantially from acute systemic inflammation in terms of tissue architecture, cellular milieu, and the sustained nature of the pathological stimulus. In this study, we evaluated the therapeutic efficacy of FKC in a human TNF (hTNF) transgenic mouse which is an extensively validated model of chronic erosive polyarthritis driven by constitutive TNF signaling [22], and recapitulates the cardinal histopathological and molecular features of human RA [23]. Using clinical RA scoring, locomotor assessment, cytokine profiling, histological analysis, and immunofluorescence of key TNFR1 downstream signaling molecules, we demonstrate that FKC rescues the arthritic phenotype in hTNF transgenic mice through inhibition of TNFR1-driven inflammatory cascade, thereby reducing synovial inflammation and joint destruction and establishing its preclinical efficacy in a mouse model of RA.

## Materials and Methods

### Peptide synthesis

FKC (FKCRRWQWRMKK-NH_2_) was synthesized using solid phase F-moc chemistry with a purity exceeding 95% (Bio Basic Inc.). The C-terminal amide modification was incorporated to protect the peptide from carboxypeptidase-mediated proteolytic degradation, thereby enhancing in vivo stability and its half-life in mice. FKC lyophilizate was reconstituted in sterile phosphate-buffered saline (PBS) (Sigma-Aldrich, D8537) at a working concentration of 5 mg/mL and passed through a 0.22 µm sterile filter immediately prior to administration.

### Animal studies

All animal procedures were approved by the Institutional Animal Care and Use Committee (IACUC) at Syracuse University under protocol number 24-004. Eight female hTNF transgenic mice of the strain B6.Cg-Tg(TNF)#Xen (tg/wt) and four female non-transgenic controls (wt/wt) were obtained at 8 weeks of age (Taconic Biosciences). We selected female mice for this study given the higher prevalence, incidence, and severity in females than males in human [24] and TNF transgenic mice [25]. All animals were housed at the animal facility at Syracuse University under pathogen-free conditions and a 12-hour light/dark cycle with access to standard rodent chow and water ad libitum. Mice were allocated to three experimental groups with respective intraperitoneal injections: (1) non-transgenic controls (PBS, N=4), (2) hTNF vehicle control (PBS, N=4), and (3) hTNF peptide treated (FKC, N=4). FKC was administered to mice at 40 mg/kg of body weight, a therapeutically effective dose which was established in our previous study [17]. Intraperitoneal injections were administered beginning at 8-week-old, once every other day, for a total of 60 days, with the experimental endpoint at 16-week-old. A blood sample from each mouse was collected via cardiac puncture into a BD Vacutainer Serum Tubes (Thermo Fisher Scientific, #12396929). Mouse spleen and joint tissues were also collected for subsequent analysis.

### Clinical arthritis scoring

Arthritis severity was assessed independently for each of the four limbs using a 0 to 3 ordinal scale, with each score corresponding to a defined clinical state based on previously established criteria [26]. The mean arthritis score per animal was computed as the average of scores across all four limbs, yielding a maximum score of 3. All scoring was performed daily by an observer blinded to the treatment group assignments throughout the duration of the experiment.

### Locomotor assessment

Hindlimb grip strength was assessed as a measure of muscular force output and hindlimb joint function using a digital grip strength meter (IMADA Co.). Locomotor activity was assessed in a standard open field maze (OFM) for 5 minutes. Each mouse was placed at the center of the arena and allowed to explore freely. Total distance traveled was measured using ANY-maze tracking software (Stoelting Co.).

### Histological analysis

Ankle joints were collected and fixed in 4% paraformaldehyde (Thermo Scientific) in PBS at 4°C for 24 hours. Following fixation, joint tissues were transferred to 14% disodium EDTA (pH 7.4) (Fisher Scientific) at 4°C for decalcification over two weeks. Decalcified tissues were cryoprotected by overnight immersion in 30% sucrose in PBS at 4°C, then embedded in Tissue-Tek O.C.T. compound (Sakura Finetek) and snap-frozen. Serial cryosections of 20 µm thickness were obtained using a cryostat and collected on Superfrost Plus glass slides (Epredia, 22-042-936). For hematoxylin and eosin (H&E) staining, tissue sections were stained with modified Mayer’s hematoxylin and eosin Y alcoholic stain (StatLab MasterTech). Stained sections were then dehydrated through a graded ethanol series, cleared in xylene, and cover slipped with Permount mounting medium (Fisher Scientific, SP15-100). For toluidine blue O (TBO) staining, cryosections were stained with 0.05% TBO solution (Spectrum Chemical) at pH 4.0 for 10 minutes and dehydrated through a graded ethanol series and cover slipped. For tartrate-resistant acid phosphatase (TRAP) staining, a TRAP staining kit (ApexBio, K2606) was applied to cryosections following manufacturer’s protocol. Brightfield images were acquired using a Leica THUNDER Imager Tissue at 20X magnification with image quantification performed by ImageJ.

### Immunofluorescence staining

Cryosections (20 µm) of the joints were permeabilized with 0.3% Triton X-100 (Bio-Rad, #1610407), then blocked with 3% bovine serum albumin (Fisher Scientific, #251856) in tris-buffered saline (J.T. Baker, #4109-02) containing 0.1% Tween-20 (TBST) (Bio-Rad, #1706531) to prevent non-specific binding. Primary antibodies were diluted 1:200 in blocking buffer and incubated with tissue sections overnight. Following primary antibody incubation, sections were washed with TBST and incubated with the appropriate species-specific Alexa Fluor (AF)- conjugated secondary antibodies at a 1:400 dilution for one hour at room temperature. Sections were subsequently washed with TBST and cover slipped using SlowFade Gold Antifade Mountant with DAPI (Invitrogen, #1063821A). Primary antibodies used include anti-TNFR1 (Abcam, ab223352); anti-TNFα (Abcam, 52B83); anti-phospho-IκBα (CST, #2859); anti-IκBα (Proteintech, 66418-1-Ig); anti-MMP3 (Proteintech, 66338-1-Ig); anti-phospho-p65 (Proteintech, 10745-1-AP); anti-p65 (CST, #8242); anti-CD68 (Abcam, ab955); and anti-CD45 (Novus Biologicals, NB100-77417). Secondary antibodies used include anti-rabbit AF488 (Abcam, ab150077), anti-mouse AF568, (Abcam, ab175473), and anti-rat AF568 (Abcam, ab175475). Fluorescence images were acquired using a Leica DMi8 inverted microscope at 25X magnification with image quantification performed by ImageJ.

### Enzyme-linked immunosorbent assay (ELISA)

Mouse plasma was extracted from collected mouse blood by centrifugation at 2,500 x g at 4°C for 15 minutes. Mouse ELISA kits, TNF (ProteinTech, KE10002), MCP-1 (ProteinTech, KE10006) IFN-γ (ProteinTech, KE10001), IL-1α (ProteinTech, KE10024), IL-1β (ProteinTech, KE10003), and IL-6 (ProteinTech, KE10007) were used to quantify cytokine levels following manufacturer’s protocols.

### Statistical analysis

All statistical analyses were performed using GraphPad Prism 11. Statistical analysis was conducted by using one-way ANOVA with post hoc Tukey or Kruskal-Wallis test for multiple comparisons. Statistical significance was defined as \**P* < 0.05, \*\**P* < 0.01, \*\*\**P* < 0.001, \*\*\*\**P* < 0.0001, and ns indicates non-significance.

## Results and Discussion

To evaluate the therapeutic efficacy of FKC against TNF-driven chronic arthritis, we performed intraperitoneal injection of FKC at 40 mg/kg into hTNF transgenic mice every other day starting at 8-week-old (day 0) and monitored disease progression until 16-week-old (day 60) (**Fig. 1B**). While control mice show no symptom of arthritis, hTNF mice administered with vehicle-only treatment exhibit a progressive increase in arthritis score to 2.6 (**Fig. 1C**) as well as swollen joints combined with contractures in mouse paws (**Fig. 1D**), recapitulating key features of human RA. Importantly, FKC treatment markedly reduces the arthritis score to 1.4 (**Fig. 1C**) and alleviates arthritis symptoms (**Fig. 1D**) in hTNF mice. The swollen joints and pain-related movement suppression are also causing the hTNF mice to have a lower hindlimb grip strength (**Fig. 1E**) and shorter distance traveled in the OFM (**Fig. 1F**) as compared to control mice. Consistent with improved joint function, FKC restores hindlimb grip strength (**Fig. 1E**), and increases the total distance traveled in the OFM (**Fig. 1F**). To determine whether functional improvement was associated with reduced systemic inflammation and decreased immune activation, we assessed splenomegaly by measuring spleen weight and quantified plasma cytokines by ELISA. Our results show that FKC significantly prevents spleen enlargement (**Fig. 1G**) and reduces cytokine levels of TNF, MCP-1, IFN-γ, IL-1α, IL-1β, and IL-6 that were elevated in hTNF mice (**Fig. 1H**), indicating an overall attenuation of systemic inflammation and immune activation.

To determine whether FKC was inhibiting TNFR1 signaling to prevent inflammation, we performed immunofluorescence staining with markers against TNFR1 and associated downstream signaling molecules in mouse joint tissues. As baseline control, the expression level of TNFR1 remains unchanged across all experimental groups (**Fig. 2A**). Downstream of TNFR1, phosphorylation of IκBα (pIκBα) was markedly elevated in hTNF mouse joints and significantly reduced with FKC treatment (**Fig. 2B**). Consistently, total IκBα, which is degraded upon pathway activation, was depleted in hTNF mouse joints and restored by FKC (**Fig. 2C**). To probe NF-κB activation, phosphorylation of p65 (pp65) was elevated in hTNF mouse joints and significantly suppressed by FKC (**Fig. 2D**), while total p65 level remains unchanged (**Fig. 2E**). Consequently, FKC treatment also decreases TNF (**Fig. 2F**) and matrix metalloproteinase-3 (MMP3) (**Fig. 2G**) levels in joint tissues, which was elevated in hTNF mice. Collectively, these findings demonstrate that FKC suppresses TNFR1-mediated inflammatory signaling in vivo by disrupting downstream signaling and inhibiting NF-κB activation, thus attenuating a NF-κB driven feed-forward inflammatory loop [27]. These results align with our previous work showing that FKC allosterically targets a conformationally active region of TNFR1, disrupts receptor dynamics, and uncouples adaptor protein recruitment without affecting receptor expression or ligand function [17]. Thus, FKC exerts its anti-inflammatory effects through selective allosteric inhibition of TNFR1, providing a mechanistic basis for its therapeutic efficacy in chronic RA.

**Figure 2.**
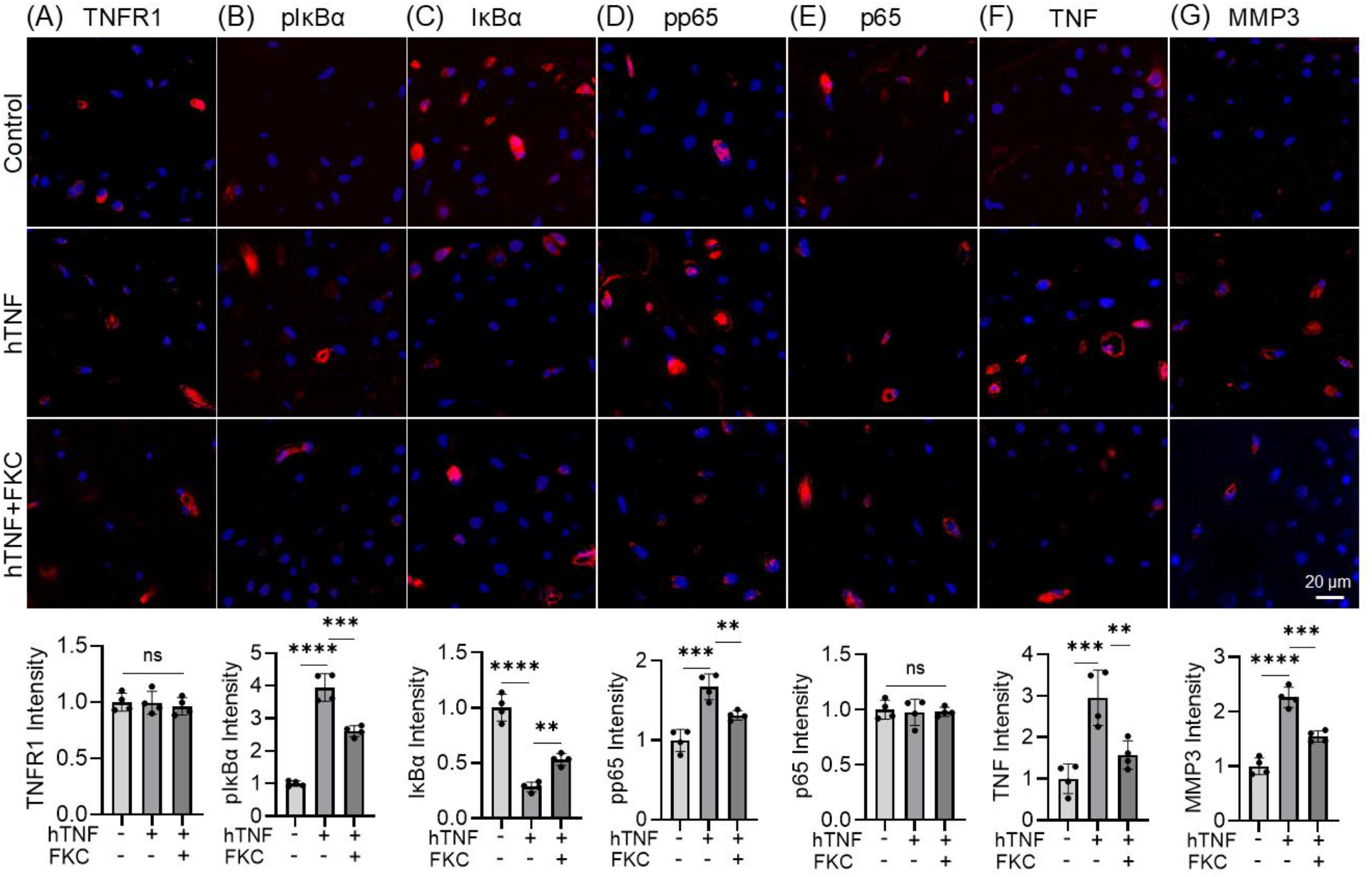
Representative immunofluorescence images and quantification of signal intensity in joint sections from control, hTNF, and hTNF+FKC mice. Sections were stained (red) for (A) TNFR1, (B) phosphorylated IκBα (pIκBα), (C) IκBα, (D) phosphorylated p65 (pp65), (E) p65, (F) TNF, and (G) matrix metalloproteinase-3 (MMP3), with DAPI nuclear counterstain (blue). hTNF mice show increased pIκBα, pp65, TNF and MMP3 levels, along with reduced total IκBα, all of which are reversed by FKC treatment. In contrast, TNFR1 and total p65 levels remain unchanged across groups. These results indicate that FKC inhibits TNFR1-mediated NF-κB activation while preserving receptor expression and NF-κB subunit abundance. Data are presented as mean ± SD (N = 4 mice). **P < 0.01, ***P < 0.001, ****P < 0.0001, ns indicates non-significance by one-way ANOVA with post hoc Tukey’s test for multiple comparisons.

To assess RA synovitis in hTNF mice, we quantified macrophage and immune cell infiltration using CD68 and CD45 immunofluorescence. Consistent with TNFR1 activation, CD68^+^ macrophages, key mediators of RA inflammation and cytokine propagation, were increased in hTNF joints and reduced by FKC treatment (**Fig. 3A**). In addition, CD45^+^ pan-leukocyte infiltration was elevated in hTNF joints and attenuated by FKC (**Fig. 3B**). To evaluate joint pathology and FKC efficacy, we performed histological analysis of hTNF mouse joints to assess synovial inflammation and the progression through cartilage degradation to terminal bone erosion, using the SMASH scoring system [23]. H&E staining displayed marked synovial hyperplasia, macrophage and immune cell infiltration, and pannus formation in hTNF mouse joints, which were attenuated by FKC. (**Fig. 3C**). Following inflammatory progression, TBO staining showed robust cartilage proteoglycan matrix in controls, marked depletion in hTNF mice, and substantial preservation with FKC treatment (**Fig. 3D**). At the terminal stage of this cascade, TRAP staining revealed increased osteoclasts at sites of bone damage in hTNF mouse joints, which were reduced by FKC (**Fig. 3E**). Collectively, these findings establish that FKC suppresses TNF/TNFR1-driven inflammatory milieu underlying RA pathology and limits progressive tissue damage in a chronic RA mouse model.

**Figure 3.**
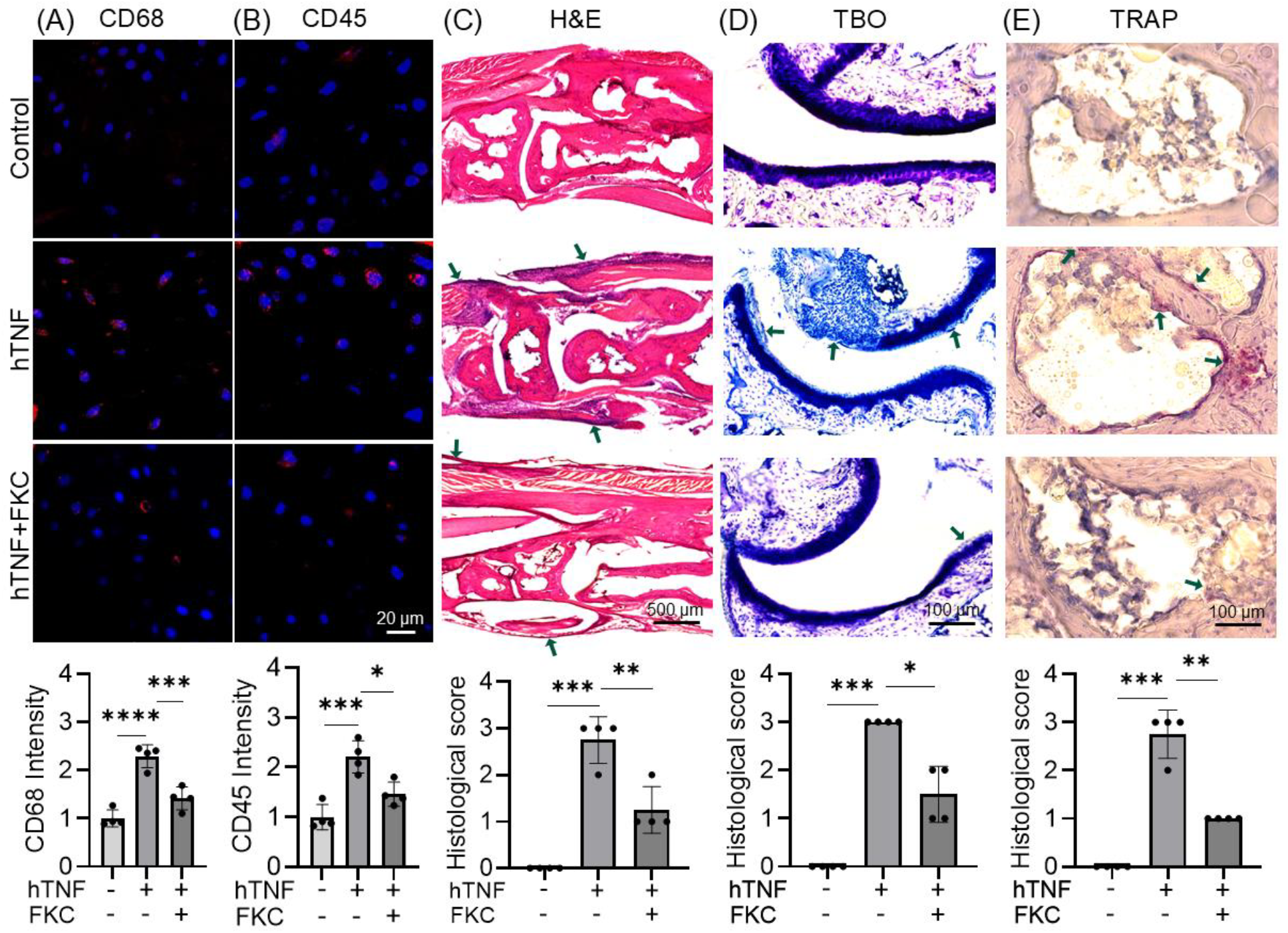
Representative immunofluorescence images and quantification of signal intensity in joint sections from control, hTNF, and hTNF+FKC mice stained for (A) the macrophage marker CD68 (red) and (B) the pan-leukocyte marker CD45 (red), with DAPI nuclear counterstain (blue). FKC treatment markedly reduces macrophage and leukocyte infiltration within the joint. (C-E) Representative histological images and corresponding quantitative scoring of joint sections across conditions. (C) Hematoxylin and eosin (H&E) staining reveals pronounced synovial hyperplasia, inflammatory cell infiltration, and pannus formation in hTNF mice, all of which are attenuated by FKC treatment. (D) Toluidine Blue (TBO) staining highlights sulfated proteoglycan content as an indicator of cartilage integrity. hTNF joints exhibit substantial proteoglycan loss, which is preserved following FKC treatment. (E) TRAP staining identifies tartrate-resistant acid phosphatase-positive osteoclasts (red-purple) at sites of bone resorption. Osteoclast activity is elevated in hTNF joints and reduced upon FKC treatment. For C-E, arrows indicate regions of inflammatory infiltration, proteoglycan loss, and osteoclast activity. Blinded semi-quantitative scoring of H&E, TBO, and TRAP sections confirms that FKC significantly reduces synovial inflammation, cartilage degradation, and osteoclast-mediated bone resorption compared to vehicle-treated hTNF mice. Data are presented as mean ± SD (N = 4 mice). For (A-B): *P < 0.05, ***P < 0.001, ****P < 0.0001 (one-way ANOVA with post hoc Tukey’s test). For (C–E): *P < 0.05, **P < 0.01, ***P < 0.001 (one-way ANOVA with post hoc Kruskal-Wallis test).

The hTNF mouse is a clinically relevant TNF/TNFR1-driven RA model that provides strong evidence of FKC target engagement. Unlike current TNF-neutralizing biologics, FKC allosterically inhibits TNFR1 signaling while preserving physiological TNF functions [17] and demonstrates efficacy in attenuating joint inflammation in vivo, supporting receptor-targeted allosteric modulation as a therapeutic strategy for RA. Although anti-TNF agents may currently appear more effective in reducing RA score of hTNF mice [28], FKC may offer greater selectivity with reduced off-target effects [17], underscoring the need to further optimize dosing, pharmacological properties, and lead peptide sequence to enhance its therapeutic potency. Given the central role of TNFR1 signaling in regulating inflammatory and immune responses, FKC is expected to similarly suppress disease progression in other RA mouse models such as collagen-induced arthritis and may have broader applicability as a potential therapeutic for autoimmune disorders in general [29,30].

## Conclusion

We demonstrate that the peptide-based allosteric TNFR1 inhibitor FKC attenuates inflammation and chronic RA pathology in hTNF mice. FKC improved clinical scores, grip strength, and locomotor function, and preserved joint architecture, effects associated with inhibition of TNF/TNFR1-driven inflammation and immune cell infiltration. Collectively, these findings establish FKC as a promising TNFR1 receptor-specific intervention for RA, support further development of allosteric TNFR1 inhibitors, and suggest potential applicability for treatment of other autoimmune disorders.

## Acknowledgements

This study is supported by a start-up fund to C.H.L. from the Department of Biology at Syracuse University and a start-up fund to J.Z. from the Department of Biomedical and Chemical Engineering at Syracuse University.

## References

1. Ma, Y.; Chen, H.; Lv, W.; Wei, S.; Zou, Y.; Li, R.; Wang, J.; She, W.; Yuan, L.; Tao, J.; et al. Global, Regional and National Burden of Rheumatoid Arthritis from 1990 to 2021, with Projections of Incidence to 2050: A Systematic and Comprehensive Analysis of the Global Burden of Disease Study 2021. Biomark. Res. 2025, 13, 47, doi:10.1186/s40364-025-00760-8.

2. Wu, D.; Huang, Y.; Zhao, J.; Long, W.; Wang, B.; Wang, Y.; Chen, H.; Wu, R. Synovial Macrophages Drive Severe Joint Destruction in Established Rheumatoid Arthritis. Sci. Rep. 2025, 15, 12111, doi:10.1038/s41598-025-93784-x.

3. Komatsu, N.; Takayanagi, H. Mechanisms of Joint Destruction in Rheumatoid Arthritis — Immune Cell–Fibroblast–Bone Interactions. Nat. Rev. Rheumatol. 2022, 18, 415–429, doi:10.1038/s41584-022-00793-5.

4. Kondo, N.; Kuroda, T.; Kobayashi, D. Cytokine Networks in the Pathogenesis of Rheumatoid Arthritis. Int. J. Mol. Sci. 2021, 22, doi:10.3390/ijms222010922.

5. Najm, A.; Ferguson, L.D.; McInnes, I.B. Cytokine Pathways Driving Diverse Tissue Pathologies in Rheumatoid Arthritis. Arthritis & Rheumatology 2026, doi:10.1002/art.43376.

6. Li, Y.; Ye, R.; Dai, H.; Lin, J.; Cheng, Y.; Zhou, Y.; Lu, Y. Exploring TNFR1: From Discovery to Targeted Therapy Development. J. Transl. Med. 2025, 23, 71, doi:10.1186/s12967-025-06122-0.

7. Taylor Peter C; Matucci Cerinic, Marco; Alten, Rieke; Avouac, Jérôme; Westhovens, Rene Managing Inadequate Response to Initial Anti-TNF Therapy in Rheumatoid Arthritis: Optimising Treatment Outcomes. Ther. Adv. Musculoskelet. Dis. 2022, 14, 1759720X221114101, doi:10.1177/1759720X221114101.

8. Siegmund, D.; Wajant, H. TNF and TNF Receptors as Therapeutic Targets for Rheumatic Diseases and Beyond. Nat. Rev. Rheumatol. 2023, 19, 576–591, doi:10.1038/s41584-023-01002-7.

9. Chen, S.; Feng, Z.; Wang, Y.; Ma, S.; Hu, Z.; Yang, P.; Chai, Y.; Xie, X. Discovery of Novel Ligands for TNF-α and TNF Receptor-1 through Structure-Based Virtual Screening and Biological Assay. J. Chem. Inf. Model. 2017, 57, 1101–1111, doi:10.1021/acs.jcim.6b00672.

10. Saddala, M.S.; Huang, H. Identification of Novel Inhibitors for TNFα, TNFR1 and TNFα-TNFR1 Complex Using Pharmacophore-Based Approaches. J. Transl. Med. 2019, 17, 215, doi:10.1186/s12967-019-1965-5.

11. Lo, Chih Hung; Vunnam, Nagamani; Lewis, Andrew K; Chiu, Ting-Lan; Brummel, Benjamin E; Schaaf, Tory M; Grant, Benjamin D; Bawaskar, Prachi; Thomas, David D; Sachs, Jonathan N An Innovative High-Throughput Screening Approach for Discovery of Small Molecules That Inhibit TNF Receptors. SLAS DISCOVERY: Advancing the Science of Drug Discovery 2017, 22, 950–961, doi:10.1177/2472555217706478.

12. Vunnam, N.; Yang, M.; Lo, C.H.; Paulson, C.; Fiers, W.D.; Huber, E.; Been, M.; Ferguson, D.M.; Sachs, J.N. Zafirlukast Is a Promising Scaffold for Selectively Inhibiting TNFR1 Signaling. ACS Bio & Med Chem Au 2023, 3, 270–282, doi:10.1021/acsbiomedchemau.2c00048.

13. Wang, X.; Guo, F.; Zhang, Y.; Wang, Z.; Wang, J.; Luo, R.; Chu, X.; Zhao, Y.; Sun, P. Dual-Targeting Inhibition of TNFR1 for Alleviating Rheumatoid Arthritis by a Novel Composite Nucleic Acid Nanodrug. Int. J. Pharm. X 2023, 5, 100162, doi:10.1016/j.ijpx.2023.100162.

14. Lo, C.H.; Schaaf, T.M.; Grant, B.D.; Lim, C.K.-W.; Bawaskar, P.; Aldrich, C.C.; Thomas, D.D.; Sachs, J.N. Noncompetitive Inhibitors of TNFR1 Probe Conformational Activation States. Sci. Signal. 2019, 12, doi:10.1126/scisignal.aav5637.

15. Murali, R.; Cheng, X.; Berezov, A.; Du, X.; Schön, A.; Freire, E.; Xu, X.; Chen, Y.H.; Greene, M.I. Disabling TNF Receptor Signaling by Induced Conformational Perturbation of Tryptophan-107. Proceedings of the National Academy of Sciences 2005, 102, 10970–10975, doi:10.1073/pnas.0504301102.

16. Lo, C.H. Targeting the Inter-Monomeric Space of TNFR1 Pre-Ligand Dimers: A Novel Binding Pocket for Allosteric Modulators. Comput. Struct. Biotechnol. J. 2025, 27, 1335–1341, doi:10.1016/j.csbj.2025.03.046.

17. Zeng, J.; Loi, G.W.Z.; Saipuljumri, E.N.; Romero Durán, M.A.; Silva-García, O.; Perez-Aguilar, J.M.; Baizabal-Aguirre, V.M.; Lo, C.H. Peptide-Based Allosteric Inhibitor Targets TNFR1 Conformationally Active Region and Disables Receptor–Ligand Signaling Complex. Proceedings of the National Academy of Sciences 2024, 121, e2308132121, doi:10.1073/pnas.2308132121.

18. Lo, C.H.; Schaaf, T.M.; Thomas, D.D.; Sachs, J.N. Fluorescence-Based TNFR1 Biosensor for Monitoring Receptor Structural and Conformational Dynamics and Discovery of Small Molecule Modulators. In The TNF Superfamily: Methods and Protocols; Bayry, J., Ed.; Springer US: New York, NY, 2021; pp. 121–137 ISBN 978-1-0716-1130-2.

19. Wenthur, C.J.; Gentry, P.R.; Mathews, T.P.; Lindsley, C.W. Drugs for Allosteric Sites on Receptors. Annu. Rev. Pharmacol. Toxicol. 2014, 54, 165–184, doi:10.1146/annurev-pharmtox-010611-134525.

20. Lo, C.H.; Huber, E.C.; Sachs, J.N. Conformational States of TNFR1 as a Molecular Switch for Receptor Function. Protein Science 2020, 29, 1401–1415, doi:10.1002/pro.3829.

21. Lo, C.H. TNF Receptors: Structure-Function Relationships and Therapeutic Targeting Strategies. Biochimica et Biophysica Acta (BBA) -Biomembranes 2025, 1867, 184394, doi:10.1016/j.bbamem.2024.184394.

22. Keffer, J.; Probert, L.; Cazlaris, H.; Georgopoulos, S.; Kaslaris, E.; Kioussis, D.; Kollias, G. Transgenic Mice Expressing Human Tumour Necrosis Factor: A Predictive Genetic Model of Arthritis. EMBO J. 1991, 10, 4025–4031, doi:10.1002/j.1460-2075.1991.tb04978.x.

23. Hayer, S.; Vervoordeldonk, M.J.; Denis, M.C.; Armaka, M.; Hoffmann, M.; Bäcklund, J.; Nandakumar, K.S.; Niederreiter, B.; Geka, C.; Fischer, A.; et al. “SMASH” Recommendations for Standardised Microscopic Arthritis Scoring of Histological Sections from Inflammatory Arthritis Animal Models. Ann. Rheum. Dis. 2021, 80, 714–726, doi:10.1136/annrheumdis-2020-219247.

24. Black, R.J.; Cross, M.; Haile, L.M.; Culbreth, G.T.; Steinmetz, J.D.; Hagins, H.; Kopec, J.A.; Brooks, P.M.; Woolf, A.D.; Ong, K.L.; et al. Global, Regional, and National Burden of Rheumatoid Arthritis, 1990-2020, and Projections to 2050: A Systematic Analysis of the Global Burden of Disease Study 2021. Lancet Rheumatol. 2023, 5, e594–e610, doi:10.1016/S2665-9913(23)00211-4.

25. Bell, R.D.; Wu, E.K.; Rudmann, C.A.; Forney, M.; Kaiser, C.R.W.; Wood, R.W.; Chakkalakal, J. V; Paris, N.D.; Klose, A.; Xiao, G.-Q.; et al. Selective Sexual Dimorphisms in Musculoskeletal and Cardiopulmonary Pathologic Manifestations and Mortality Incidence in the Tumor Necrosis Factor–Transgenic Mouse Model of Rheumatoid Arthritis. Arthritis & Rheumatology 2019, 71, 1512–1523, doi:10.1002/art.40903.

26. Ishiwatari-Ogata, C.; Kyuuma, M.; Ogata, H.; Yamakawa, M.; Iwata, K.; Ochi, M.; Hori, M.; Miyata, N.; Fujii, Y. Ozoralizumab, a Humanized Anti-TNFα NANOBODY® Compound Exhibits Efficacy Not Only at the Onset of Arthritis in a Human TNF Transgenic Mouse but Also During Secondary Failure of Administration of an Anti-TNFα IgG. Front. Immunol. 2022, Volume 13-2022, doi:10.3389/fimmu.2022.853008.

27. Hoffmann, A.; Cheng, G.; Baltimore, D. NF-KB: Master Regulator of Cellular Responses in Health and Disease. Immunity & Inflammation 2025, 1, 2, doi:10.1007/s44466-025-00014-0.

28. Ubah, O.C.; Steven, J.; Porter, A.J.; Barelle, C.J. An Anti-HTNF-α Variable New Antigen Receptor Format Demonstrates Superior in Vivo Preclinical Efficacy to Humira® in a Transgenic Mouse Autoimmune Polyarthritis Disease Model. Front. Immunol. 2019, Volume 10-2019, doi:10.3389/fimmu.2019.00526.

29. Lo, C.H.; Zeng, J. TNF as a Mediator of Metabolic Inflammation and Body-Brain Interaction in Obesity-Driven Neuroinflammation and Neurodegeneration. Ageing Res. Rev. 2025, 112, 102891, doi:10.1016/j.arr.2025.102891.

30. Li, Y.; Ye, R.; Dai, H.; Lin, J.; Cheng, Y.; Zhou, Y.; Lu, Y. Exploring TNFR1: From Discovery to Targeted Therapy Development. J. Transl. Med. 2025, 23, 71, doi:10.1186/s12967-025-06122-0.

